# Population size shapes trade-off dilution and adaptation to a marginal niche unconstrained by sympatric habitual conditions

**DOI:** 10.1101/2023.11.13.566869

**Authors:** Yashraj Chavhan, Sarthak Malusare, Sutirth Dey

## Abstract

Niche expansion remains significantly understudied in sympatric scenarios where marginal and habitual niches are simultaneously available. Devoid of spatial constraints, such conditions impose selection to maintain fitness in habitual (high-productivity) niches while adapting to marginal (low-productivity) niches. Hence, habitual niche composition should constrain adaptation to marginal niches. This remains untested. Similarly, it is unknown if larger populations expand their niches better. We tested these hypotheses using experimental evolution with *Escherichia coli* and found that larger populations consistently adapted better to both marginal and habitual niches. Whereas the habitual niche composition (constant versus fluctuating environments; environmental fluctuations varying in both predictability and speed) significantly shaped fitness in habitual niches, surprisingly, it failed to constrain adaptation to the marginal niche. Curiously, two negatively correlated habitual niches can still each be positively correlated with the marginal niche. This allows the marginal niche to dilute trade-offs between habitual niches, thereby enabling costless niche expansion.

## Introduction

Marginal niches, where most individuals of the species under consideration have initially poor survival and reproduction, are key sources of ecological opportunities for evolutionary changes (Kawecki 2008). Adaptation to such niches can lead to significant changes in the species’ ecological capabilities, ultimately reshaping its niche width, i.e., the set of conditions that support its growth and reproduction (Hutchinson 1961; Holt and Gaines 1992). Adaptation to marginal niches has been conventionally studied in macroscopic organisms near the physical boundaries of species ranges (Hoffmann and Blows 1994), making dispersal dynamics an important determinant of this phenomenon (Kawecki 2000; Lenormand 2002). However, adaptation to marginal niches has received relatively less attention in sympatric ecological scenarios where spatial constraints are absent and the habitual niche is simultaneously available. In this study, we investigate how various population genetic and ecological factors interact to shape adaptation in bacterial populations to a new marginal niche in the presence of a habitual niche.

Consider a scenario wherein a marginal niche becomes available to an asexual bacterial population residing in an environment that simultaneously offers access to the habitual niche. In terms of nutrient niches, the mammalian gut can be taken as a representative example of such an environment (Scanlan 2019). Here a change in the host’s diet can make a novel low productivity carbon source (the marginal niche) available to the bacteria in addition to the habitual niche comprising high productivity carbon source(s) (Phillips 2009; Payne et al. 2012). Due to the marginal niche’s low productivity, bacterial growth will initially be largely dependent on the high productivity habitual niche until there is sufficient adaptation to the marginal carbon source. The first step towards such an adaptation will be the appearance of a mutation that increases the usability of the marginal carbon source. Such a mutation will then need to survive genetic drift in order for its fate to be governed by selection (Sniegowski and Gerrish 2010). The per generation rate at which mutations with a beneficial effect size *s* survive drift is *NU_b_s*, where *N* is the population size and *U_b_* is the rate of beneficial mutations (Desai and Fisher 2007; Desai et al. 2007). Consequently, larger populations (*i.e.*, with higher *N*) should adapt better to the marginal niche. Moreover, apart from having better access to rare large effect beneficial mutations, larger populations also show greater efficiency of natural selection (Desai and Fisher 2007; Neher 2013; Chavhan et al. 2019). Taken together, larger populations are expected to adapt better to the marginal niche.

An important factor that should be instrumental in shaping the dynamics of adaptation to a marginal niche is the identity of the habitual sympatric niche. This is because the selection coefficient *s* in the expression *NU_b_s* is influenced by the overall fitness in the simultaneous presence of the habitual (high productivity) and the marginal (low productivity) niches. Therefore, mutations that increase fitness in the marginal niche would have greater chance of surviving drift if they were not too deleterious in the habitual niche. Thus, maintaining fitness in the sympatric habitual niche should be an important constraint while adapting to the marginal niche. Hence, adaptation to the marginal niche should be influenced by the habitual niche’s identity. This can happen in several ways. For example, the demand to maintain fitness in the habitual niche can be considerably different in cases where the habitual niche applies a single constant selection pressure versus cases where it imposes multiple dynamically changing (fluctuating) selection pressures. Whereas fluctuating environments allow evolution to be shaped by fitness trade-offs across several individual selection pressures, evolution in constant environments that present a single principal selection pressure is oblivious to such trade-offs (Kassen 2002; Bono et al. 2017). Consequently, evolution can proceed relatively more freely in such constant environments (Bono et al. 2017). Moreover, the relative significance of fitness trade-offs in constant versus fluctuating environments has recently been shown to be dependent on the population size (Chavhan et al. 2021).

Interestingly, temporal environmental fluctuations can themselves vary vastly in terms of their speed and predictability, which can be important in shaping the ensuing selection pressures within the habitual niche. For instance, a recent experimental evolution study has shown that the speed of fluctuations (i.e., the rate of switch between environments) can be instrumental in shaping adaptive outcomes at the level of both phenotypes and genotypes (Boyer et al. 2021). Such differences in the speed of environmental fluctuations can even change the direction of selection (Salignon et al. 2018). Moreover, if the environment fluctuates predictably, microbes can evolve adaptive strategies to anticipate future changes (Tagkopoulos et al. 2008). However, such anticipation is unlikely if the fluctuations are unpredictable (Karve et al. 2015, 2016). Agreeing with these notions, several empirical studies have suggested that the predictability of environmental fluctuations should be an important determinant of evolutionary trajectories (Mitchell et al. 2009; Dhar et al. 2013). Finally, multiple experimental evolution studies have shown that the predictability of environmental fluctuations significantly shapes evolutionary outcomes (Hughes et al. 2007; Boyer et al. 2021 but see Karve et al. 2018). Taken together, the habitual niche can apply a variety of selection pressures depending on whether it remains constant or fluctuates over time. Furthermore, the selection pressures within the habitual niche should be shaped by the speed and predictability of the corresponding environmental fluctuations. Such diversity of selection pressures from the habitual niche should lead to diverse fitness correlations between the habitual niche and the marginal niches. To our knowledge, the effects of all these factors (in combination with the population size) and their interactions on adaptation to marginal niches remain uninvestigated.

We conducted experimental evolution with *E. coli* at two population sizes in several constant and fluctuating environments to determine the population genetic and ecological determinants of the adaptation to a marginal niche under sympatric availability of habitual niches. We also investigated if and how population size interacted with the habitual niche’s identity to shape fitness in high productivity habitual niches. We further determined if fitness in fluctuating habitual niches is shaped by the interactions of population size and the speed and predictability of environmental fluctuations. To our knowledge, such three-way interactions have not been put to experimental tests yet. As expected, larger populations consistently adapted better to the marginal niche regardless of the habitual niche’s identity. Surprisingly, we found that, despite the differences in the constant and fluctuating selection pressures applied by the various habitual niches across our treatments, adaptation to the marginal niche was not influenced by the identity and stability of the sympatric habitual niche(s). Larger populations consistently gained greater fitness in both constant and fluctuating habitual niches. Interestingly, the identities of selection pressures within constant habitual niches were key to fitness changes in them. However, neither the speed nor the predictability of environmental fluctuations significantly influenced adaptation to the fluctuating habitual niches. Remarkably, our study revealed the occurrence of non-transitive fitness correlations across three distinct niches. Incorporating this non-transitivity within an extension of Fisher’s Geometric Model accounted for our observations across both the habitual and marginal niches in both constant and fluctuating environments at multiple population sizes. Our results elucidate substantial evolutionary potential for rapid and costless expansion of bacterial niche width in the face of a new sympatric ecological opportunity, even at low mutational supply rates.

## Materials and Methods

### Experimental evolution

We founded *E. coli* MG1655 populations from a single common ancestral clone and cultured them in eight different environmental conditions at two distinct population sizes (Fig. 1; see Appendix S1 for detailed information about the ancestral strain and culture media). All the eight environments offered acetate as the low productivity carbon source that constituted the marginal niche. Out of the eight environments, four offered a single distinct high productivity carbon source (constant habitual niche) for bacterial growth (one of thymidine (Thy), galactose (gal), sorbitol (Sor) or arabinose (Ara)). In the other four environments, the habitual niche (high productivity carbon source) fluctuated over time in all combinations of predictable versus unpredictable and fast (switching every ∼13.3 generations) versus slow (switching every ∼40 generations) fluctuations. We designated these fluctuations in the habitual niche as PF (predictable and fast), PS (predictable and slow), UF (unpredictable and fast) and US (unpredictable and slow). Taken together, a combination of two different population sizes and eight different environments gave rise to 16 distinct evolutionary regimens (Thy-L, Thy-S, Gal-L, Gal-S, Sor-L, Sor-S, Ara-L, Ara-S, PFL, PFS, PSL, PSS, UFL, UFS, USL and USS; here L and S refer to large and small population sizes respectively; Fig. 1). We propagated six independently evolving biological replicates of each regimen, making a total of 96 independently evolving experimental populations. All the populations were allowed to evolve for ∼480 generations in 96-well plates at a culture volume of 300 µl incubated at 37° C. We followed the standard methodology for growing microbial populations of different sizes at identical culture volumes (Desai et al. 2007; Raynes et al. 2014; Vogwill et al. 2016). The large (L) populations underwent a 1:10 periodic bottleneck after every 12 hours. The ten-fold growth between successive bottlenecks corresponded to ∼3.3 generations. In contrast, the small (S) populations were bottlenecked 1:10^4^ every 48 hours. The ten-thousand-fold growth between successive bottlenecks in these populations corresponded to ∼13.3 generations. This bottlenecking protocol ensured that for a given environmental regimen, both the large and the small populations spent similar durations in the stationary phase.

**Fig. 1.**
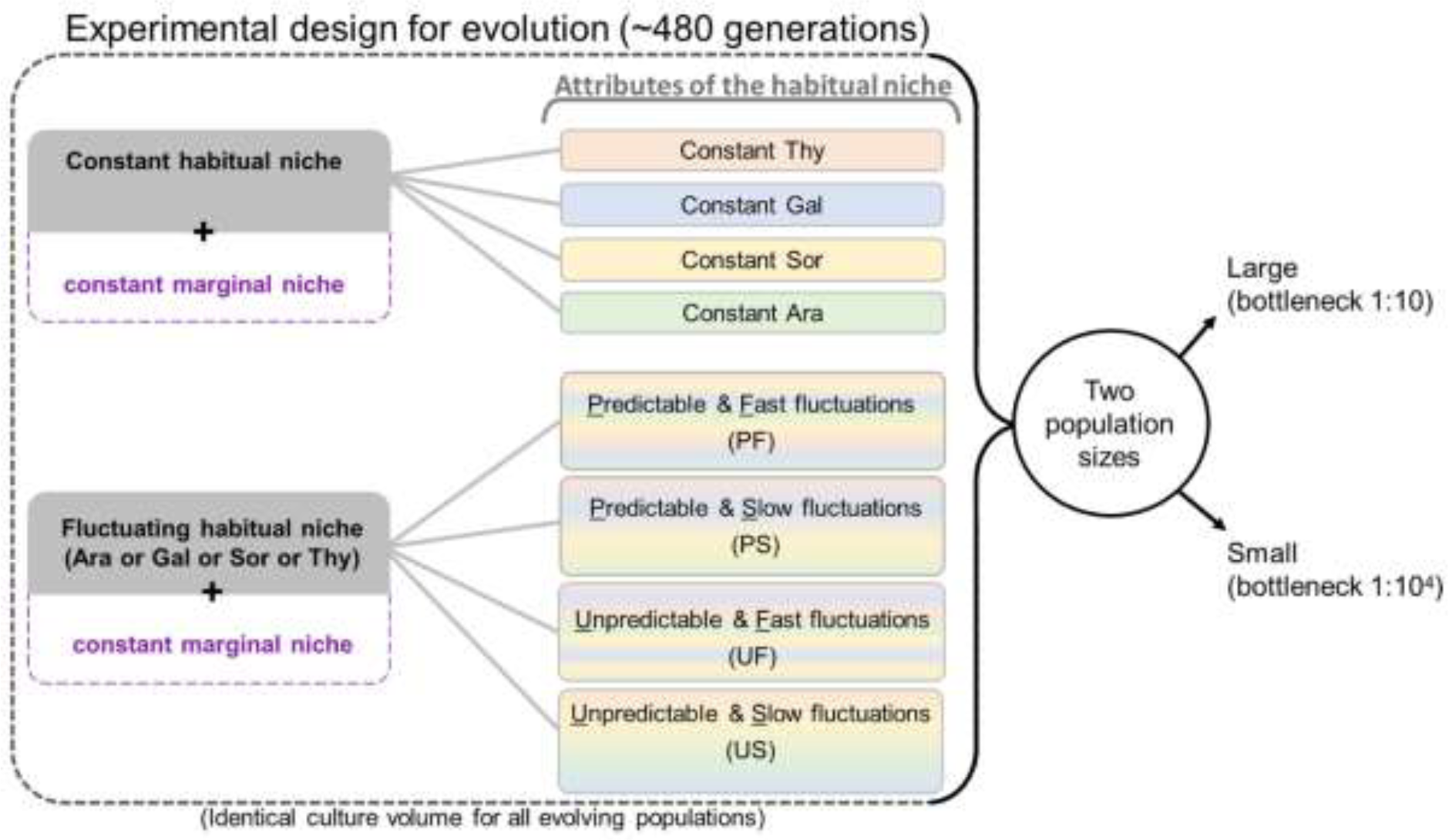
A schematic representation of the experimental design used in this study. There were sixteen distinct environmental regimens (populations derived from a common ancestor evolved in eight different environmental conditions at two different population sizes). In all the eight environmental conditions, the marginal niche (acetate) was always present in addition to the habitual niche, which had distinct attributes in different environments. See the text for details.

### Fitness measurement

At the end of the experiment, we measured the fitness of each of the 96 independently evolving populations in five different environments (where the sole carbon source was one of acetate, Thy, Gal, Sor or Ara). To this end, we first revived samples derived from the endpoint cryo-stocks by allowing them to grow hundred-fold in a glucose-based M9 minimal medium. We then grew all the 96 revived populations in each of the five environments and measured their optical density at 600 nm every 20 minutes using an automated well-plate reader (Synergy HT, BIOTEK^®^ Winooski, VT, USA). The physical conditions during growth measurement were identical to those during the evolution experiment. As a single 96 well-plate was insufficient for all the fitness assays, we used a randomized complete block design (RCBD) and assayed one replicate of each of the sixteen regimens in each environment on a given day (Milliken and Johnson 2009). We used the maximum slope of the growth curves, calculated over a moving window of ten readings as the measure of fitness (Leiby and Marx 2014; Karve et al. 2015; Chavhan et al. 2021).

### Statistical analysis

We determined if our experimental populations had adapted or maladapted significantly to their habitual and marginal niches. To this end, we used single-sample t-tests against the ancestral fitness (scaled to 1 in each of the five assay environments) and corrected for family-wise error rates using the Holm–Šidàk procedure, categorizing fitness > 1 (corrected *P* < 0.05) as adaptations and cases with fitness < 1 (corrected *P* < 0.05) as maladaptations. For the eight regimens with fluctuating habitual niches, we used the geometric means of the fitness values in Thy, Gal, Sor and Ara (the habitual niche components) and compared them to the ancestral value (=1) using single sample t-tests followed by Holm–Šidàk correction.

To investigate the effects of population size and the habitual niche’s identity on fitness in the marginal niche, we used a mixed model ANOVA (RCBD) with ‘population size’ (two levels: L and S) and ‘habitual niche’ (eight levels: Ara, Gal, Sor, Thy, PF, PS, UF and US) as fixed factors crossed with each other and ‘day of assay’ as the random factor. We also used partial *η*^2^ as the measure of effect size (Cohen 1988), where small, medium and large effects were identified with partial *η*^2^ < 0.06, 0.06 < partial *η*^2^ < 0.14, and 0.14 < partial *η*^2^, respectively.

We used a similar mixed model ANOVA (RCBD) to analyse how population size and the identity of the constant habitual niche shaped fitness in the latter. Specifically, we treated ‘population size’ (two levels: L and S) and ‘constant habitual niche identity’ (four levels: Thy, Gal, Ara and Sor) as fixed factors crossed with each other and ‘day of assay’ as the random factor.

Analogously, to analyse how population size and the fluctuating niche’s identity shaped fitness in the latter, we conducted a mixed model ANOVA (RCBD) on geometric mean fitness values over the four component environments of the fluctuating habitual niche. Here we treated ‘population size’ (two levels: L and S), ‘fluctuation predictability’ (two levels: predictable and unpredictable) and ‘fluctuation speed’ (two levels: fast and slow) as fixed factors crossed with each other and ‘day of assay’ as the random factor.

## Results

### Adaptation to the marginal niche was shaped by the population size, not by the habitual niche’s identity or stability

Our evolution experiment resulted in widespread adaptation to exploit the marginal ecological opportunity presented by Acetate. Specifically, fifteen out of the 16 regimens underwent significant adaptation to acetate during the ∼480 generations of evolution (Fig. 2; see Table S1 for statistical details). Moreover, fifteen out of the 16 regimes avoided significant declines in fitness in their habitual niches. Interestingly, twelve out of the 16 regimens also adapted significantly to their habitual niches (Tables S2 and S3). Next, we determined the population genetic and ecological factors that could explain the extent of adaptation to the marginal niche.

**Fig. 2.**
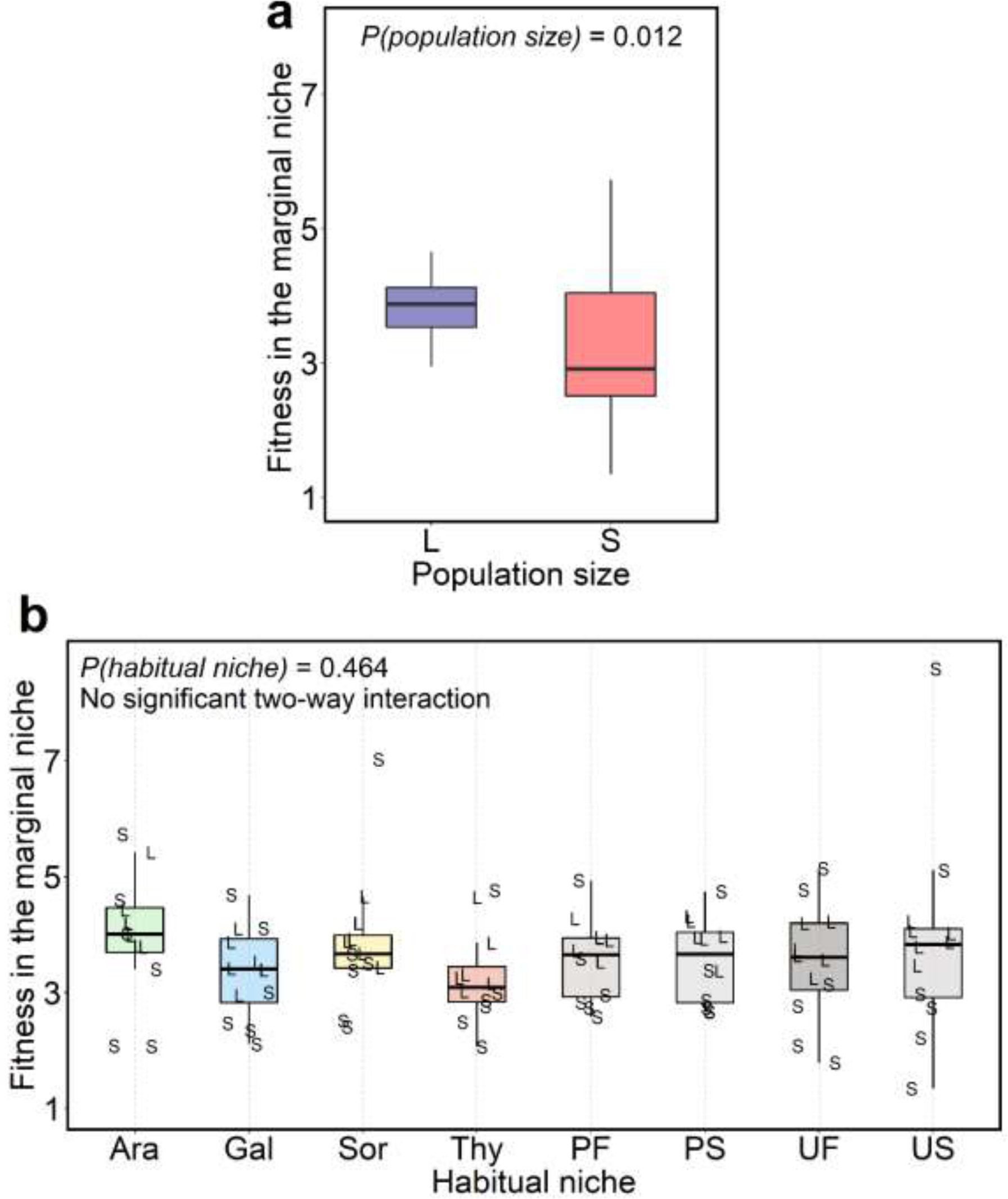
Fitness in the marginal niche after ∼480 generations. **(a)** Larger populations had significantly better fitness in the marginal niche as compared to smaller populations. **(b)** The habitual niche’s identity did not have a significant influence on fitness in the marginal niche. The colored plots correspond to regimens with a constant habitual niche, black and white plots correspond to regimens with a fluctuating habitual niche. L and S represent large and small populations, respectively. In both the plots, the lower and upper hinges of boxes show the 25% and 75% quantiles, respectively. The horizontal line within boxes represents the median. The whiskers denote the two extremes within the data that are less than or equal to 1.5 times the length of the box extending away from the box. While the larger populations adapted better to the marginal niche, surprisingly, such adaptation was unconstrained by the composition of the habitual niche.

We found that the population size had a significant effect on fitness in the marginal niche (mixed model ANOVA: Population size (fixed effect) *F_1,75_* = 6.62; *P* = 0.01; partial η^2^ = 0.081 (medium effect)). Specifically, larger populations gained higher fitness in acetate (Fig. 2a). This observation can be explained by the relatively greater supply of variation and better efficiency of natural selection in larger populations (Desai and Fisher 2007; Desai et al. 2007; Chavhan et al. 2019, 2020).

Unexpectedly, the identity of the habitual niche (constant (Ara, Gal, Sor, or Thy) or fluctuating (PF, PS, UF, US)) did not have a significant effect on fitness in the marginal niche (*F_7,75_* = 0.96; *P* = 0.46) (Fig. 2b). Moreover, we did not find any significant statistical interaction of the population size and the habitual niche’s identity in shaping the fitness in the marginal niche (*F_7,75_* = 0.20; *P* = 0.98).

This absence of significant effects of the habitual niche’s identity on adaptation in the marginal niche is surprising due to several reasons: Firstly, acetate could only support severely restricted growth in the ancestor (Fig. S1). Consequently, most of the growth in our experimental regimens depended on the habitual niche, particularly during the initial phases of evolution. Hence, all the sixteen regimens faced a strong selection to maintain (or gain) fitness in their respective habitual niches. Indeed, twelve out of 16 regimens adapted to their respective habitual niches while three out of 16 regimens maintained their fitness at the ancestral level in them (Table S3). Secondly, while the constant habitual niches of the Ara-L, Ara-S, Gal-L, Gal-S, Thy-L, and Thy-S regimens offered a single nutritionally challenging selection pressure, the fluctuating habitual environment niches in the PFL, PFS, PSL, PSS, UFL, UFS, USL, and USS regimens offered multiple dynamic selection pressures. Hence, the selection pressure(s) were expected to be different in constant versus fluctuating habitual niches. Since acetate was the only stable component in the environments faced by regimes with fluctuating habitual niches, significant adaptation to acetate is expected in these regimens despite the differences in niche fluctuations across them. Thirdly, two of the individual carbon sources used in the habitual niches in our experiments (Gal and Thy) are expected to impose selection in markedly different directions. Specifically, in a previous study with *E. coli*, we had shown that fitness in Gal is negatively correlated with fitness in Thy (Chavhan et al. 2020). Indeed, we found that the Gal-L and Gal-S regimens adapted to Gal but became maladapted to Thy, a carbon source that they had never encountered during our experiment (Table S2). Likewise, Thy-L gained significant fitness in Thy, but maladapted to Gal. Interestingly, we also found similar trends with fitness changes in Sor. Sor-L adapted to Sor but lost significant fitness in Gal and Thy. Analogously, both Gal-L and Gal-S lost significant fitness in Sor (Table S2). Thus, the selection pressures posed by the individual habitual carbon sources used here are expected to be different. In contrast to the above pattern of costs in regimes with constant habitual niches, the dynamic environment of the PFL, PSL, PSS, UFL, UFS, USL, and USS regimens fluctuating across Ara, Gal, Sor and Thy made them evade all fitness pairwise fitness costs among these four habitual carbon sources (Table S2; PFS paid a marginally significant cost across Gal and Sor). Finally, multiple empirical investigations have shown that differences in the speed and predictability of environmental fluctuations can take populations towards significantly different evolutionary fates (Hughes et al. 2007; Boyer et al. 2021). Thus, differences in terms of predictability and speed of fluctuations in the habitual niches of the PFL, PFS, PSL, PSS, UFL, UFS, USL, and USS regimes are expected to influence their adaptive trajectories. Taken together, the selection pressures imposed by the habitual niche were not only instrumental in shaping the evolution in our experiment, but they were also expected to be markedly different across different regimens. Since the correlated effects of different selection pressures are also expected to be different, it is surprising that the identity of the habitual niche did not affect fitness in the marginal niche.

Before presenting a population genetic explanation of this unexpected pattern of niche expansion and discussing its implications, we describe the fitness changes in the habitual niches of our experimental regimens. Since the constant habitual niches in our study were qualitatively different from the fluctuating habitual niches, we conducted separate analyses of the drivers of adaptation in them.

### Fitness gains in constant habitual niches were shaped by their identities and the population size

We found that larger populations tended to have significantly greater fitness in their constant habitual niches (mixed model ANOVA: Population size (fixed effect) *F_1,35_* = 8.099; *P* = 0.007; partial η^2^ = 0.188 (large effect)) (Fig. 3). In contrast to fitness in the marginal niche, the identities of the habitual niches significantly affected fitness changes in them (mixed model ANOVA: identity of the constant habitual niche (fixed effect) *F_3,35_* = 136.08; *P* < 10^−5^; partial η^2^ = 0.921 (large effect)) (Fig. 3). There was no statistically significant interaction of population size and constant habitual niche’s identity in influencing fitness in the habitual niche (*F_3,35_* = 1.532; *P* = 0.223). Specifically, the extent of adaptation to habitual niches showed the following trend: Thy > Gal > Ara ≈ Sor (based on post-hoc tests using Tukey’s HSD (Fig. 3; Table S2;). As described above, the individual habitual carbon sources used in our study are expected to apply markedly different selection pressures. Moreover, microbial experimental evolution studies usually show an inverse relationship between the current absolute fitness in an environment and the adaptability to that environment (the rule of declining adaptability described by Couce and Tenaillon (2015)). Indeed, we found that absolute ancestral fitness in habitual niches had an inverse trend relative to the extent of adaptation in them (Fig. S2). Thus, the rule of declining adaptability can account for the differences in the extents of adaptation to the different constant habitual niches in our experiments, thereby explaining why the identities of the constant habitual niches should significantly influence fitness gains in them (Fig. 3).

**Fig. 3.**
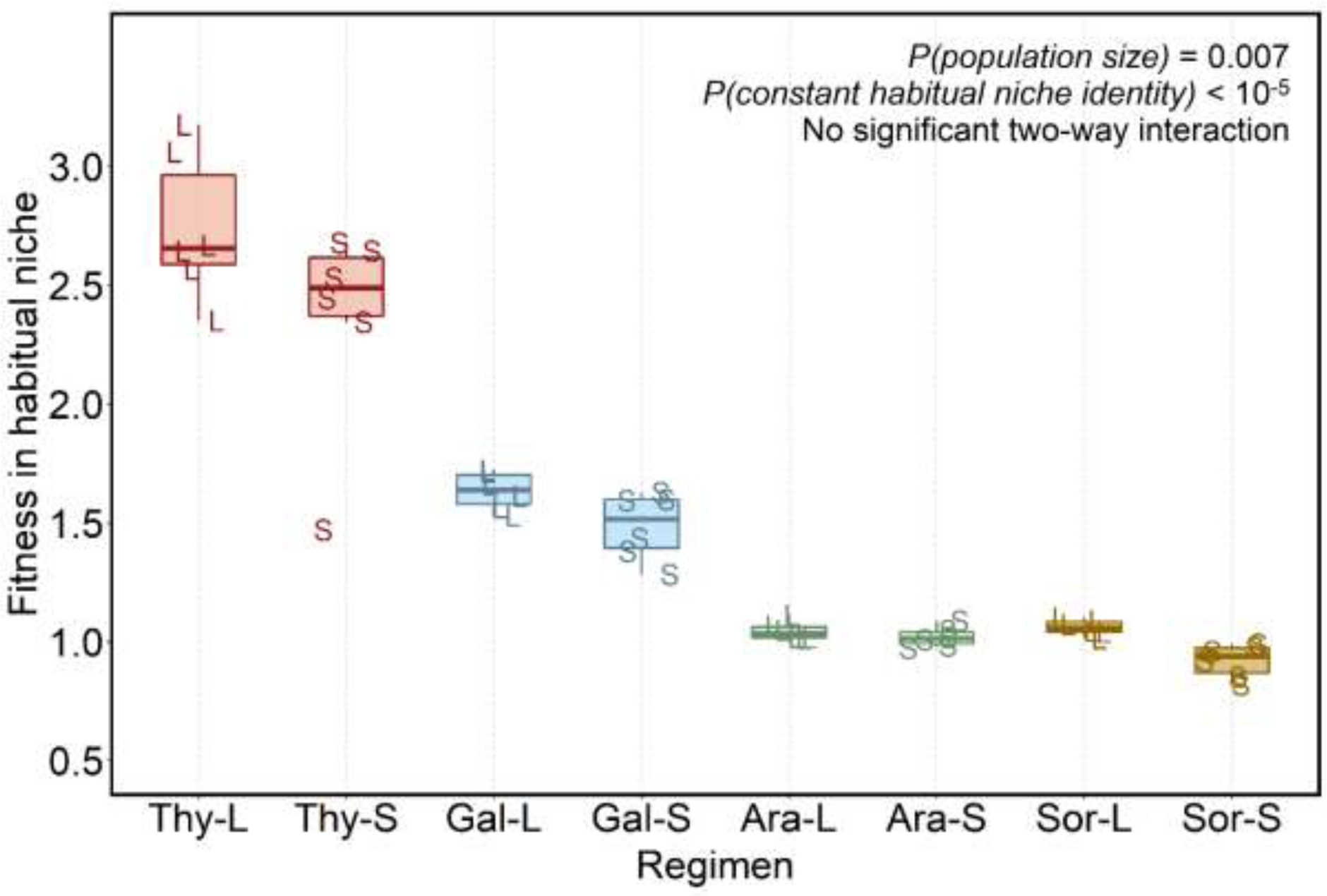
Fitness in the constant habitual niche after ∼480 generations. L and S represent large and small populations, respectively. The box and whisker notations are similar to those in Fig. 2. Overall, fitness in the constant habitual niches was influenced significantly by both the population size and the constant habitual niche’s identity, but the effects of either of these factors was not contingent on the other.

### Adaptation to the fluctuating habitual niche was not affected by the predictability or speed of fluctuations

Analogous to the observation in the constant habitual niche (Fig. 3), larger populations adapted better to their fluctuating habitual niche (Fig. 4). Specifically, we found that larger populations gained greater geometric mean fitness across Ara, Gal, Sor and Thy (mixed model ANOVA: *F_1,35_* =104.43; *P* = 4.792 × 10^−12^). Surprisingly, the predictability (*F_1,35_* =0.003; *P* = 0.954) and speed (*F_1,35_* = 0.526; *P* = 0.473) of fluctuations failed to significantly shape fitness changes in the fluctuating habitual niche (Fig. 4). This trend was so unambiguously evident that neither the three possible two-way interactions nor the three-way interaction between the three fixed effects were significant: population size × predictability (*F_1,35_* =1.191; *P* = 0.283); population size × speed (*F_1,35_* = 0.409; *P* = 0.527); predictability × speed (*F_1,35_* = 0.230; *P* = 0.635); population size × predictability × speed (*F_1,35_* = 4.9 × 10^−4^; *P* = 0.983).

**Fig. 4.**
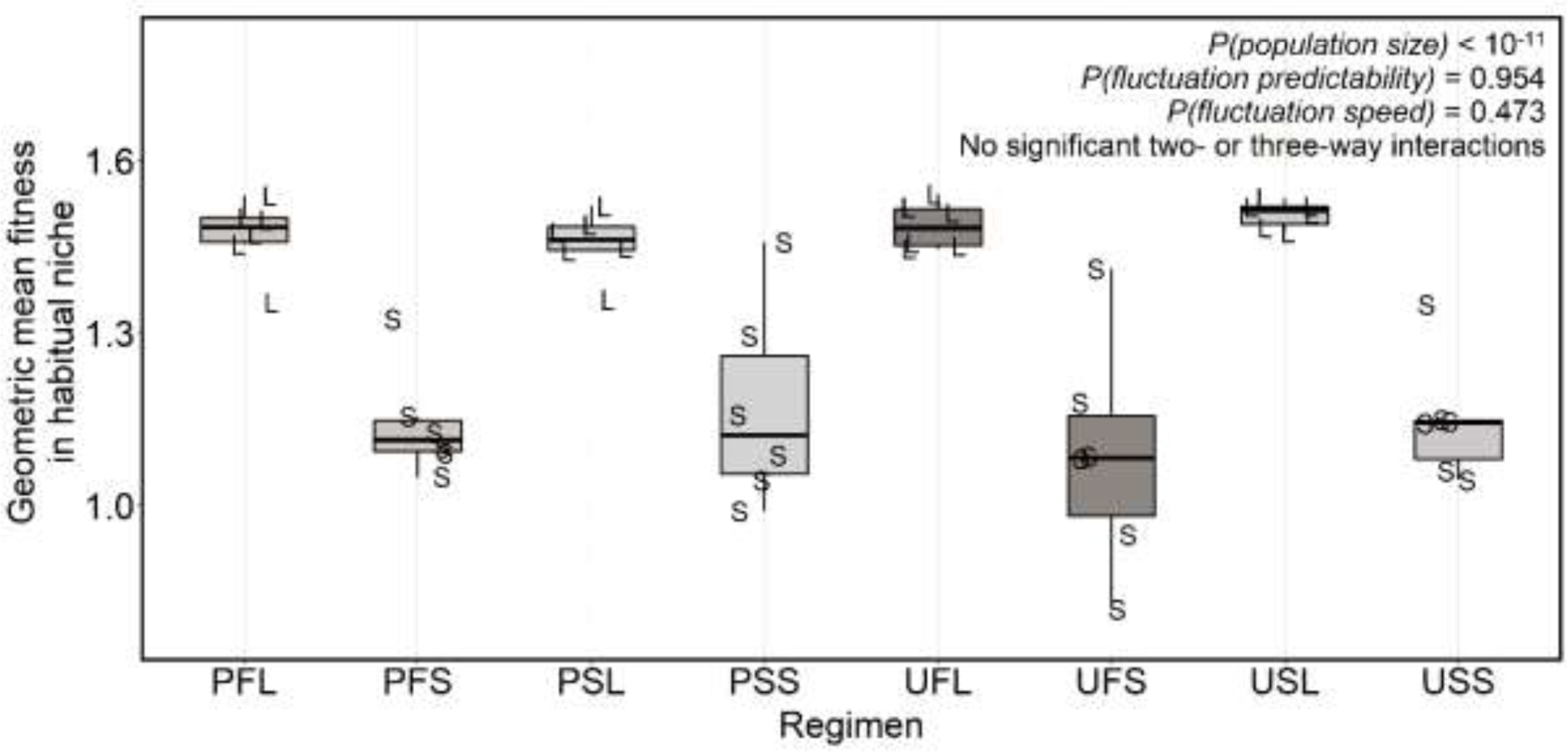
Fitness in the fluctuating habitual niche after ∼480 generations. L and S represent large and small populations, respectively. The box and whisker notations are similar to those in Fig. 2. Although the fitness in fluctuating habitual niches was significantly shaped by the population size, neither the predictability nor the speed of habitual niche fluctuations was instrumental to it.

### Non-transitive fitness correlations: Two negatively correlated habitual niches can each have positive correlations with the marginal niche

Surprisingly, we found a non-transitivity in the fitness correlations across Gal, Thy, and Acetate (Fig. 5). We observed a negative Gal-Thy fitness correlation in regimens with either Gal or Thy in their constant habitual niche (Fig. 5a), which agrees with previous results Chavhan et al. (2020). However, the regimens with Gal as the habitual carbon source (Gal-L and Gal-S) showed a significant positive fitness correlation across Gal and Acetate (Fig. 5b). Analogously, Thy-L and Thy-S exhibited a significant positive correlation in fitness across Thy and Acetate (Fig. 5c).

**Figure 5.**
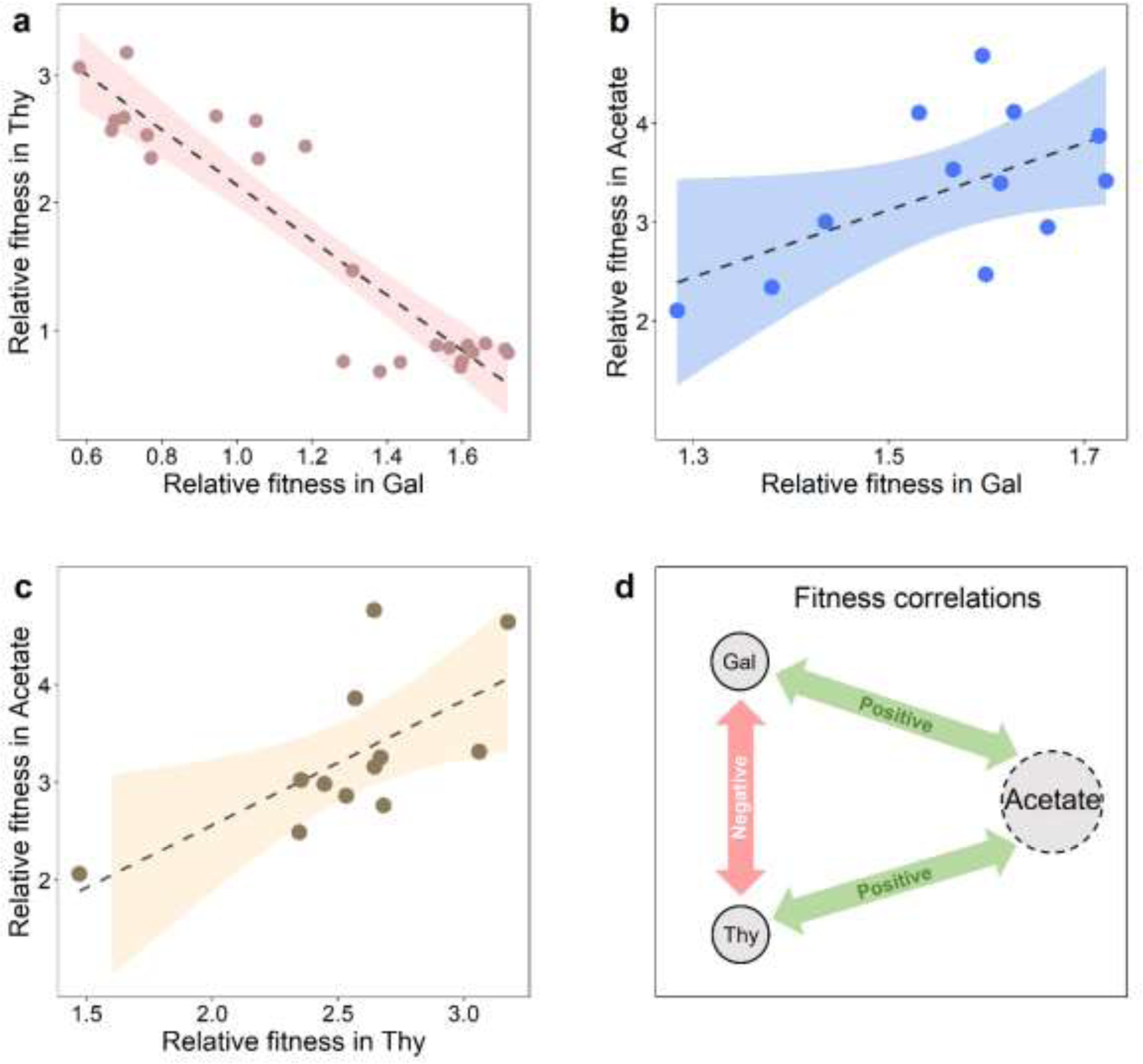
Fitness correlations across Gal, Thy and Acetate. **(a)** Fitness was negatively correlated across Gal and Thy in regimens that had either Gal or Thy as the only carbon source in their constant habitual niche (Spearman’s ρ = −0.767; *P* = 1.231 × 10^−5^). **(b)** There was a positive correlation across Gal and Acetate in regimens that had Gal as the only carbon source in their habitual niche (Spearman’s ρ = 0.573; *P* = 0.051). **(c)** Fitness was significantly positively correlated across Thy and Acetate in regimens that had Thy as the only carbon source in their habitual niche (Spearman’s ρ = 0.670; *P* = 0.017). **(d)** Taken together, our experiment revealed a non-transitivity in fitness correlations across Gal, Thy and Acetate. Thus, two habitual niche components that are negatively correlated in terms of fitness can both be positively correlated with the marginal niche. In (a), (b) and (c), the dashed line represents the best fit linear curve, and the shaded region represents 95% confidence interval of the linear regression. Taken together, we found a non-transitivity in fitness correlations across a three distinct niches (two habitual and one marginal).

Taken together, our experiment reveals rampant adaptation to a marginal niche that becomes available in addition to the sympatric habitual niche. While our results validate the expectation that adaptation to the marginal niche should be influenced by the population size, surprisingly, they also reveal that such adaptation can be consistently unconstrained by the composition or stability of the sympatric habitual niche. We now turn to the ecological and population genetic explanations of the fitness trends observed here.

## Discussion

We found that bacterial populations can quickly adapt to a new and low productivity (marginal) niche without losing fitness in a variety of sympatric habitual niches. This observation reveals a potential for widespread bacterial niche expansion in the face of novel ecological opportunities. Interestingly, such rampant adaptation to the marginal niche is a major expectation of the ‘rule of declining adaptability’ (Couce and Tenaillon 2015). Specifically, this rule predicts that our experimental populations should adapt much more to Acetate than the habitual carbon sources, to which the bacteria were already relatively more adapted to. In other words, the fitness gains in the marginal niche should be much greater than fitness changes in the habitual niches. Our observations match this general expectation (compare Fig. 2b with Fig. 3, noting the difference of the scale on the Y-axis).

In contrast to the above result, our key observation is surprising: We found that adaptation to the marginal niche was unconstrained by the identity and composition of a variety of constant and fluctuating sympatric habitual niches. When we had previously made *E. coli* evolve in constant environments containing either Gal or Thy as the only available carbon source, significant adaptation to one of the two environments always led to significant maladaptation to the other (Chavhan et al. 2020). Hence, Gal and Thy were expected to pose opposing selection pressures on our experimental populations. This was the primary reason why we had incorporated Gal and Thy within the sympatric habitual niches in the present study (which employed the same bacterial strain as Chavhan et al. (2020)). We expected that such contrasting selection pressures within the habitual niches of our different treatments would make the adaptation to the marginal niche significantly constrained by the composition of the sympatric habitual niche (which varied across eight distinct types, four with constant environments and four with fluctuating environments).

However, we found that two habitual niche components that have negative fitness correlations with each other could still show a positive correlation with the marginal niche (Fig. 5d). This can explain why the adaptation to Acetate was unconstrained by the presence of Gal or Thy in the habitual niche. More generally, one can use Fisher’s geometric model for multiple fitness optima to understand adaptation to a marginal niche in the presence of a sympatric habitual niche (Martin and Lenormand 2015; Schick et al. 2015). Due to its very nature, the marginal niche will have its fitness optimum at a large distance away from the ancestral genotype. Theory predicts that a mutation beneficial in the marginal niche would be more likely to be neutral (or beneficial) than being deleterious in the other niches (Martin and Lenormand 2015). Agreeing with this expectation, Acetate did not show a negative fitness correlation with any of the four components of the habitual niches in our study (fitness in Sor showed a weak positive correlation, and that with Ara was uncorrelated (Fig. S3)). Such absence of significant negative fitness correlations can explain why adaptation to the marginal niche was not influenced by the identity of the habitual niche.

Interestingly, the ever present Acetate diluted the Gal-Thy trade-offs due to the non-transitivity in fitness correlations discussed above. As expected based on (Chavhan et al. (2020), we found a negative Gal-Thy fitness correlation in regimens with either Gal or Thy in their constant habitual niche here (Fig. 5a). However, unlike Chavhan et al. (2020) (which employed Gal and Thy but lacked the marginal niche), adaptation to one of Gal or Thy did not always lead to maladaptation to the other. Specifically, while both Thy-L and Thy-S adapted significantly to Thy, only Thy-L became maladapted to Gal; the fitness of Thy-S in Gal remained unchanged (Table S2).

Although the Gal-Thy trade-off was diluted by Acetate, Gal and Thy still showed a negative fitness correlation (Fig. 5a). Thus, even if both Gal and Thy were present as parts of the fluctuating habitual niche, it was unlikely that a mutation could enhance fitness in both of them. Hence, simultaneous adaptation to Gal and Thy, if it happens at all, should require the enrichment of multiple distinct mutations. Since such an event is expected to be relatively rare, simultaneous adaptation to Gal and Thy should occur primarily in larger populations with high enough mutational supply for enriching multiple mutations, which is exactly what was observed in a previous experimental evolution study (Chavhan et al. (2021). Although Acetate was not a part of the nutrient media in that study, it had employed environments that fluctuated across several sole carbon sources (including Gal and Thy). In Chavhan et al. (2021), the large populations could adapt to both Gal and Thy by fixing multiple distinct mutations, but the small populations could not do so. Agreeing with these previous observations, here we found that all the large population regimens with a fluctuating habitual niche (PFL, PSL, UFL, USL) became adapted to both Gal and Thy but the analogous small population regimens (PFS, PSS, UFS, USS) could adapt to neither (Table S2).

Finally, another important observation of our study is that adaptation to the fluctuating habitual niche was not significantly shaped by the predictability or speed of fluctuations (Fig. 4). This contrasts with the results of some previous studies that have found the predictability and/or the speed of environmental fluctuations to be important determinants of fitness changes (Hughes et al. 2007; Boyer et al. 2021). Interestingly, a recent study has found that the predictability of environmental fluctuations does not significantly influence the extent of adaptation in *E. coli* (Karve et al. 2018). We note that a key difference between these previous studies and our experiment is the sustained presence of a marginal niche that the bacteria could adapt to. It is likely that adaptation to this unchanging marginal niche could have masked the nuanced effects of the speed and predictability of habitual niche fluctuations, particularly given that Acetate diluted the Gal-Thy trade-offs. Indeed, we found another manifestation of trade-off dilution in the presence of the marginal niche in the fluctuating habitual niche regimens. In one of our previous studies that did not involve Acetate, the small populations evolving in an environment that fluctuated across several sole carbon sources (including Gal and Thy) adapted to Thy but became significantly maladapted to Gal (Chavhan et al. 2021). Contrastingly, in the present study (where Acetate was always available), all the small populations evolving in the fluctuating environment regimens (PFS, PSS, UFS, USS) avoided maladaptation to Gal.

### Implications

Our observations imply a pervasive evolutionary potential for the expansion of bacterial niche width that is unconstrained by the composition of the habitual niche, even at low population sizes. Furthermore, despite employing four different habitual niches with different unchanging (constant) sole carbon sources and four others with different fluctuations across them, only one out of the 16 regimens showed fitness trade-offs across their habitual and marginal niches (Table S4). Such pervasive and costless manner of niche expansion in the face of a new ecological opportunity can potentially explain why *E. coli* tends to show a phenomenal metabolic niche breadth (Sajed et al. 2016). We note that most of the sixteen regimens in our study adapted to both the marginal and the habitual niches (Table S4). This contrasts with the results of an experimental evolution study with *Pseudomonas fluorescens* where the populations adapted largely to the low-productivity niche but not to the high productivity niche (Jasmin and Kassen 2007).

To our knowledge, this is the first study to propose that fitness trade-offs between two habitual environments can be diluted by the presence of a marginal ecological opportunity. We show that a non-transitivity in fitness correlations can lead to significant dilution of trade-offs. This finding can be an important step in understanding ecological specialization in bacteria.

Our explanations are likely to work generally in the presence of a marginal niche with a large scope of adaptation. Moreover, unlike Gal and Thy, if the environments in the habitual niche do not show reciprocal trade-offs, relatively greater niche expansion is expected. Given the relatively short span of our study (∼480 generations), our results demonstrate that bacteria can quickly adapt to multiple environmental components to expand their niches if a new opportunity becomes available, regardless of properties of the habitual niche still at their disposal. Our observations and their population genetic explanations should act as stepping-stones for more nuanced tests of niche expansion theories by targeting direct and pleiotropic mutational effects in large versus small populations in constant versus fluctuating habitual niches.

## Acknowledgements

We thank Milind Watve and M.S. Madhusudhan for their valuable inputs. YC was supported by a Senior Research Fellowship initially sponsored by IISER Pune and then by Council for Scientific and Industrial Research (CSIR), Govt. of India. YC also acknowledges Department of Biotechnology, Govt. of India for postdoctoral funding for this work. YC was supported by a postdoctoral fellowship awarded by the Wenner-Gren Foundations (Sweden) during the composition and writing of this manuscript. SM was supported by an INSPIRE undergraduate fellowship, sponsored by Department of Science and Technology (DST), Govt. of India. This project was supported by an external grant (BT/PR22328/BRB/10/1569/2016) from Department of Biotechnology, Govt. of India, and internal funding from IISER Pune.

## Supplementary Information

### Appendix SA1

#### Details of the ancestral strain and nutrient media

**Ancestral strain:** *Escherichia coli* MG1655 lacY::kan. (resistant to kanamycin).

**Nutrient media:** Our study employed four constant and four fluctuating nutrient environments. Each constant environment contained an M9-based minimal medium, 1 litre of which had:

- 12.8 g Na_2_HPO_4_.7H_2_O
- 3.0 g KH_2_PO_4_
- 0.5 g NaCl
- 1.0 g NH_4_Cl
- 240.6 mg MgSO_4_
- 11.1 mg CaCl_2_
- 4g of the pre-decided habitual carbon source
- 4g of sodium acetate (the ever-present marginal niche)
- 50 mg Kanamycin sulphate

The four constant environments differed in terms of the identity of the pre-decided habitual carbon source. The following four carbon sources were used in our experiment:

- Arabinose (Ara)
- Galactose (Gal)
- Sorbitol (Sor)
- Thymidine (Thy)

In the four fluctuating nutrient environments, the habitual carbon source fluctuated between the above four carbon sources every ∼13.3 generations while sodium acetate was constantly present as the marginal carbon source.

**Fig. S1.**
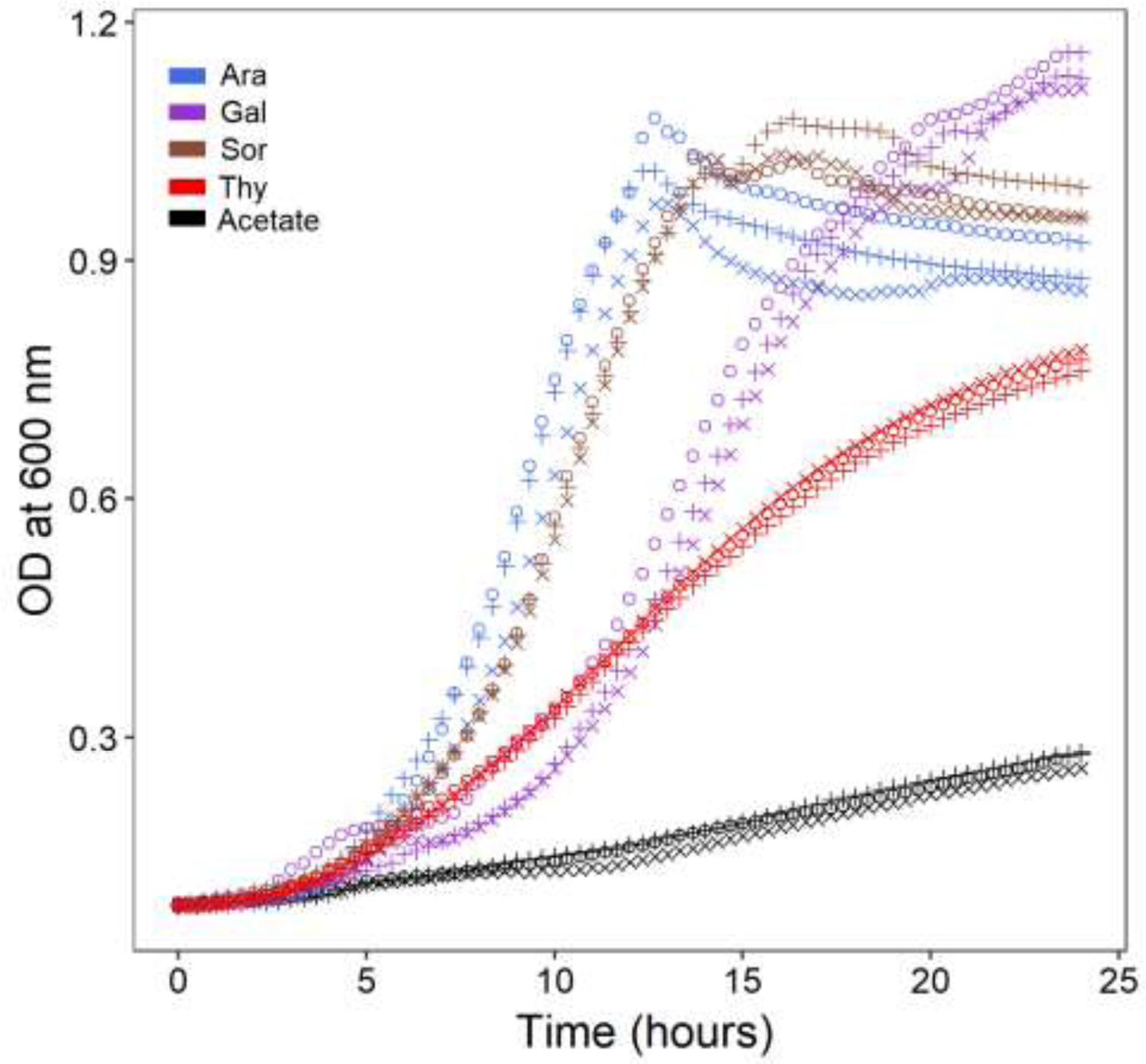
The rationale behind choosing Acetate as the marginal niche for our experiment. Acetate supported substantially lower growth than the four habitual carbon sources (Ara, Gal, Sor, and Thy). Also see Fig. S2.

**Table S1.** Fitness changes in the marginal niche single sample t-tests (*N* = 6) against the ancestral fitness (scaled to 1)). The *P* values were corrected for family-wise error rates using the Holm-Šidák procedure.

| Regimen | Mean relative fitness | Corrected P | Inference |
| --- | --- | --- | --- |
| AL | 4.270242 | 0.000237844 | Adaptation |
| AS | 3.648791 | 0.012651868 | Adaptation |
| GL | 3.544164 | 8.27514E-05 | Adaptation |
| GS | 3.1204 | 0.012781838 | Adaptation |
| SL | 3.961356 | 5.13908E-05 | Adaptation |
| SS | 3.750789 | 0.030927201 | Adaptation |
| TL | 3.541009 | 0.000461687 | Adaptation |
| TS | 2.987307 | 0.013380837 | Adaptation |
| PFL | 3.887881 | 5.22008E-06 | Adaptation |
| PFS | 3.269059 | 0.00606959 | Adaptation |
| PSL | 3.963065 | 1.09478E-05 | Adaptation |
| PSS | 3.176525 | 0.006217328 | Adaptation |
| UFL | 3.767876 | 3.00656E-05 | Adaptation |
| UFS | 3.280626 | 0.039815349 | Adaptation |
| USL | 3.905231 | 3.76782E-06 | Adaptation |
| USS | 3.827156 | 0.176057133 | No change |

**Table S2.** Adaptation or maladaptation to the habitual carbon sources (single sample t-tests (*N* = 6) against the ancestral fitness (scaled to 1)). The *P* values were corrected for family-wise error rates using the Holm-Šidák procedure.

| Regimen | Assay Environment | Mean | Corrected $P$ | Inference |
| --- | --- | --- | --- | --- |
| Ara-L | Ara | 1.047095 | 0.09061878 | No change |
| Ara-L | Gal | 0.899821 | 0.118017773 | No change |
| Ara-L | Sor | 0.939063 | 0.091235169 | No change |
| Ara-L | Thy | 1.092703 | 0.091377083 | No change |
| Ara-S | Ara | 1.020637 | 0.289682695 | No change |
| Ara-S | Gal | 0.743547 | 0.000546261 | Maladaptation |
| Ara-S | Sor | 0.913148 | 0.011743599 | Maladaptation |
| Ara-S | Thy | 0.836784 | 0.008789995 | Maladaptation |
| Gal-L | Ara | 1.083268 | 0.036119868 | Adaptation |
| Gal-L | Gal | 1.635322 | 2.99278E-05 | Adaptation |
| Gal-L | Sor | 0.941378 | 0.011327888 | Maladaptation |
| Gal-L | Thy | 0.868418 | 0.000197829 | Maladaptation |
| Gal-S | Ara | 0.966397 | 0.257217716 | No change |
| Gal-S | Gal | 1.487345 | 0.001513162 | Adaptation |
| Gal-S | Sor | 0.874405 | 0.018835832 | Maladaptation |
| Gal-S | Thy | 0.749709 | 0.000345593 | Maladaptation |
| Sor-L | Ara | 0.976053 | 0.587810092 | No change |
| Sor-L | Gal | 0.767935 | 0.001536821 | Maladaptation |
| Sor-L | Sor | 1.061002 | 0.012506628 | Adaptation |
| Sor-L | Thy | 0.779143 | 5.33282E-05 | Maladaptation |
| Sor-S | Ara | 0.843587 | 0.063991603 | No change |
| Sor-S | Gal | 0.687172 | 0.001714285 | Maladaptation |
| Sor-S | Sor | 0.919832 | 0.037892971 | Maladaptation |

Table S2 (continued)
| Regimen | Assay Environment | Mean | Corrected P | Inference |
| --- | --- | --- | --- | --- |
| Sor-S | Thy | 0.774939 | 0.010935145 | Maladaptation |
| Thy-L | Ara | 1.029067 | 0.12302482 | No change |
| Thy-L | Gal | 0.683121 | 0.000235651 | Maladaptation |
| Thy-L | Sor | 0.918037 | 0.212261581 | No change |
| Thy-L | Thy | 2.745439 | 0.000186149 | Adaptation |
| Thy-S | Ara | 1.024005 | 0.049883592 | Adaptation |
| Thy-S | Gal | 1.050872 | 0.539227194 | No change |
| Thy-S | Sor | 0.786947 | 0.052396836 | MS* Maladaptation |
| Thy-S | Thy | 2.353409 | 0.003588368 | Adaptation |
| PFL | Ara | 1.037809 | 0.321088381 | No change |
| PFL | Gal | 1.630182 | 5.12645E-05 | Adaptation |
| PFL | Sor | 1.018063 | 0.467792143 | No change |
| PFL | Thy | 2.714323 | 7.17729E-06 | Adaptation |
| PFS | Ara | 1.023830 | 0.227554022 | No change |
| PFS** | Gal | 1.521371 | 0.050918783 | MS* Adaptation |
| PFS | Sor | 0.933590 | 0.000929809 | Maladaptation |
| PFS | Thy | 1.336395 | 0.318635345 | No change |
| PSL | Ara | 1.036134 | 0.292249186 | No change |
| PSL | Gal | 1.637865 | 3.03757E-06 | Adaptation |
| PSL | Sor | 1.011076 | 0.669998756 | No change |
| PSL | Thy | 2.619420 | 8.85578E-07 | Adaptation |
| PSS | Ara | 1.023830 | 0.905874664 | No change |
| PSS | Gal | 1.521371 | 0.100675218 | No change |

Table S2 (continued)
| Regimen | Assay Environment | Mean | Corrected P | Inference |
| --- | --- | --- | --- | --- |
| PSS | Sor | 0.933590 | 0.156892634 | No change |
| PSS | Thy | 1.336395 | 0.284669337 | No change |
| UFL | Ara | 1.073203 | 0.00503885 | Adaptation |
| UFL | Gal | 1.613365 | 2.09643E-06 | Adaptation |
| UFL | Sor | 1.051203 | 0.008350401 | Adaptation |
| UFL | Thy | 2.691810 | 5.58899E-06 | Adaptation |
| UFS | Ara | 1.008975 | 0.750354776 | No change |
| UFS | Gal | 1.211113 | 0.575771079 | No change |
| UFS | Sor | 0.938349 | 0.044188397 | Maladaptation |
| UFS | Thy | 1.408203 | 0.514402368 | No change |
| USL | Ara | 1.072171 | 0.002246683 | Adaptation |
| USL | Gal | 1.656833 | 7.28375E-07 | Adaptation |
| USL | Sor | 1.036104 | 0.0441302 | Adaptation |
| USL | Thy | 2.805214 | 5.4264E-07 | Adaptation |
| USS | Ara | 0.976209 | 0.594372297 | No change |
| USS | Gal | 1.087440 | 0.625291022 | No change |
| USS | Sor | 0.944795 | 0.187145628 | No change |
| USS | Thy | 1.972183 | 0.11481443 | No change |

\*Marginally significant

**Table S3.** Changes in the geometric mean fitness in the eight regimens that evolved in fluctuating habitual niches. The geometric mean fitness was calculated across the four components of the fluctuating habitual niches (Ara, Gal, Sor, and Thy) and analysed using single sample t-tests against the ancestral level (scaled to 1).

| Regimen | Geometric mean fitness | *P* | Inference |
| --- | --- | --- | --- |
| PFL | 1.469528 | 0.000010 | Adaptation to the fluctuating habitual niche |
| PSL | 1.455601 | 0.000006 | Adaptation to the fluctuating habitual niche |
| UFL | 1.487332 | 0.000001 | Adaptation to the fluctuating habitual niche |
| USL | 1.507005 | 0.000000 | Adaptation to the fluctuating habitual niche |
| PFS | 1.141479 | 0.015476 | Adaptation to the fluctuating habitual niche |
| PSS | 1.171889 | 0.061628 | Marginally significant adaptation to the fluctuating habitual niche |
| UFS | 1.087625 | 0.337133 | No change |
| USS | 1.148681 | 0.020324 | Adaptation to the fluctuating habitual niche |

**Fig. S2.**
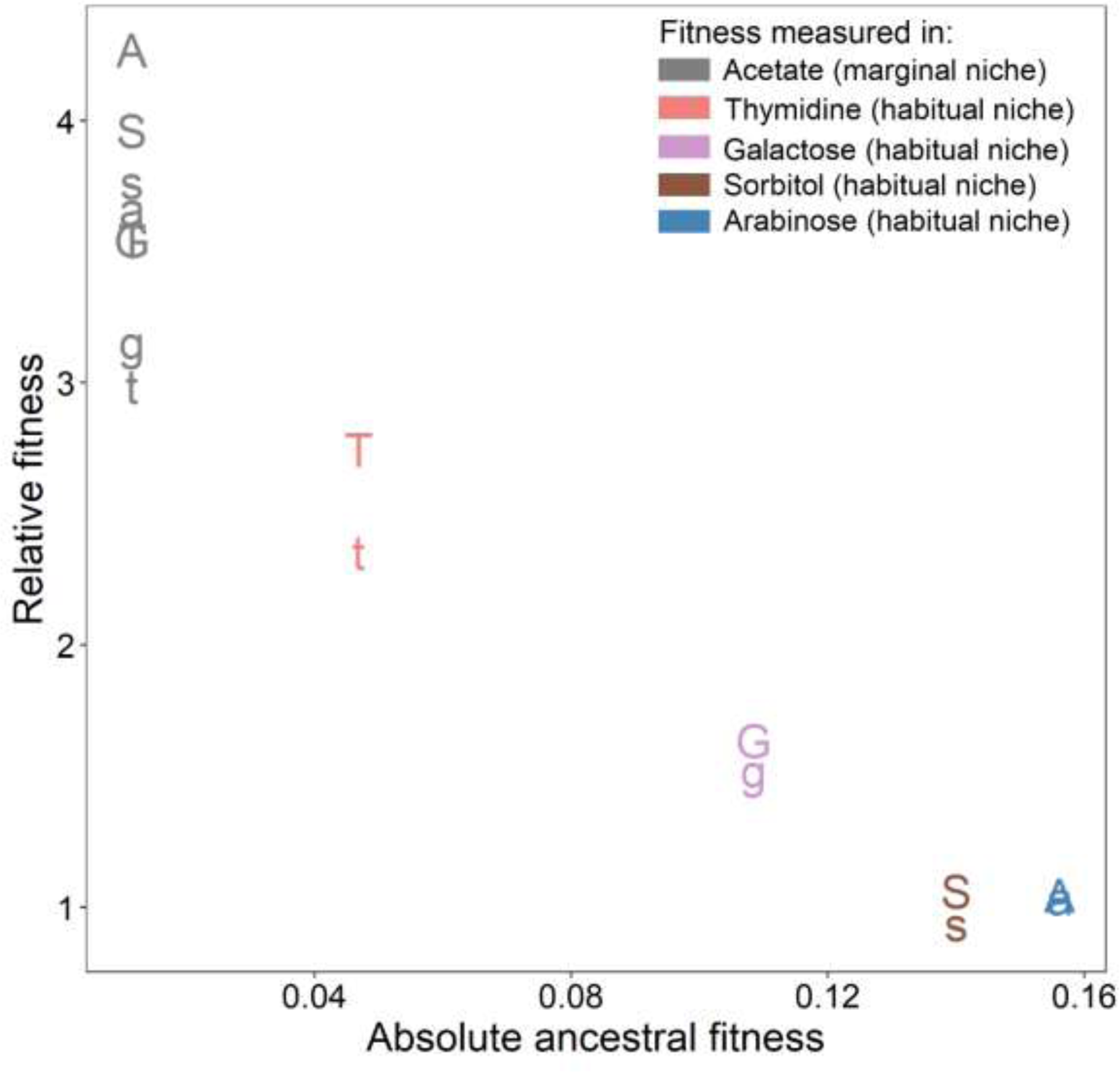
Our experiment aligned with the expectations of the “rule of declining adaptability.” Across the five distinct nutritional niches in our study (one marginal (Acetate) and four habitual (Ara, Gal, Sor, and Thy), the absolute ancestral fitness showed an inverse relationship with the relative fitness gained by regimens with unchanging habitual niches. Mean relative fitness values are plotted (N = 6), A and a refer to Ara-L and Ara-S, respectively. Similarly, G represent Gal-L while g represents Gal-S; S stands for Sor-L while s refers to Sor-S; T refers to Thy-L, t stands for Thy-S.

**Fig. S3.**
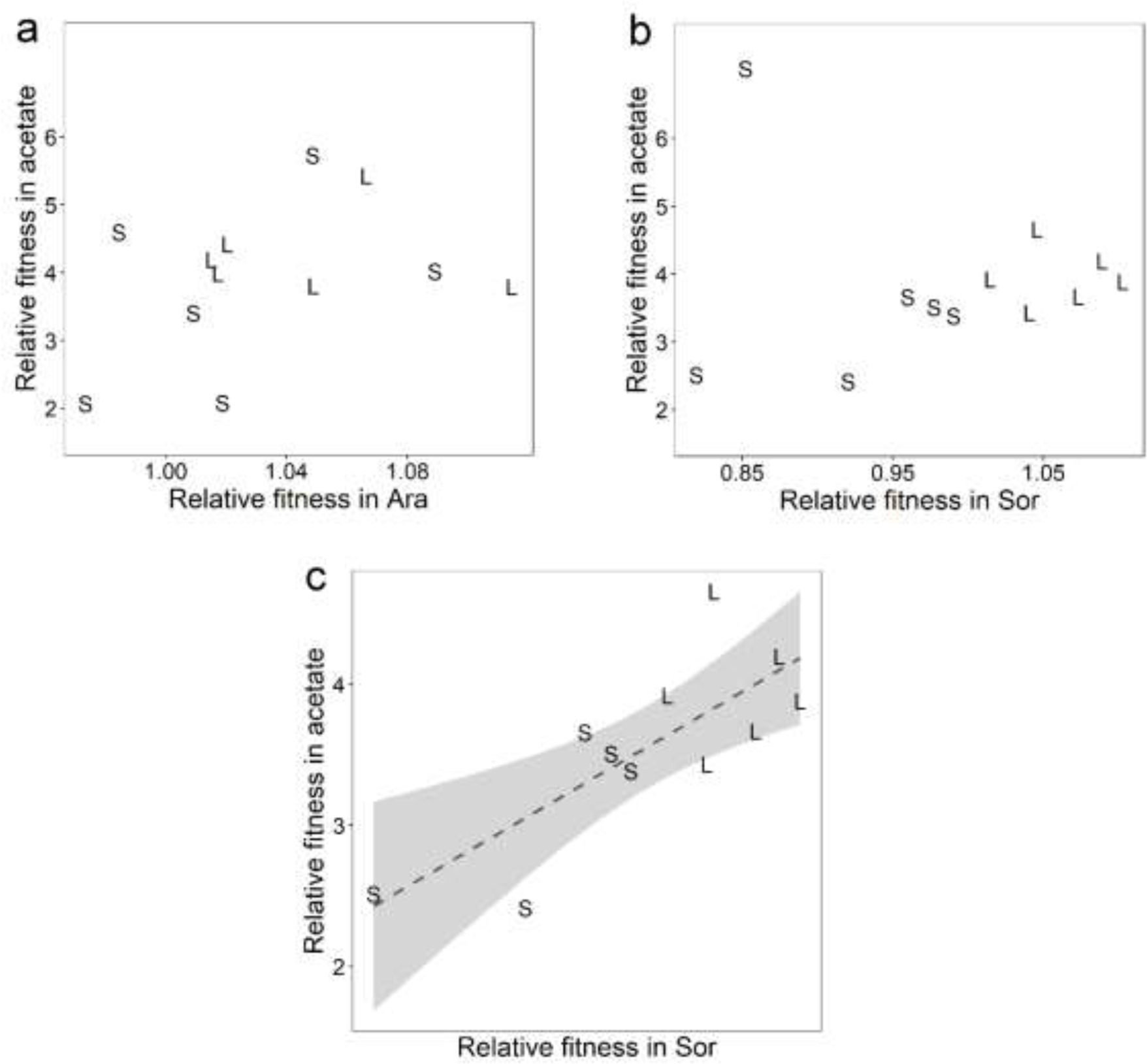
Fitness in Acetate was uncorrelated with that in Ara while Acetate and Sor showed a weakly positive fitness correlation. L and S refer to large and small population regimens, respectively. **(a)** There was no significant fitness correlation across Acetate and Ara in regimens that had Ara as the only carbon source in their habitual niche (Ara-L and Ara-S): Spearman’s ρ = 0.252; *P* = 0.430. **(b)** Among regimes that had Sor as the only habitual carbon source (Sor-L and Sor-S), the fitness correlation across Acetate and Sor was not significant when all the 12 data points were included (Spearman’s ρ = 0.413; *P* = 0.183). **(c)** Upon excluding the outlier with relative fitness in Acetate > 6, the fitness correlation across Acetate and Sor in Sor-L and Sor-S was positive (Spearman’s ρ = 0.745; *P* = 0.008).

**Table S4.** Fitness trade-offs across the habitual and marginal niches in our experimental regimens.

| **Regimen** | **Fitness change in the marginal niche** | **Fitness change in the habitual niche** | **Fitness trade-off across the marginal and habitual niches** |
| --- | --- | --- | --- |
| **Ara-L** | Adaptation | No change | No |
| **Ara-S** | Adaptation | No change | No |
| **Gal-L** | Adaptation | Adaptation | No |
| **Gal-S** | Adaptation | Adaptation | No |
| **Sor-L** | Adaptation | Adaptation | No |
| **Sor-S** | Adaptation | Maladaptation | Yes |
| **Thy-L** | Adaptation | Adaptation | No |
| **Thy-S** | Adaptation | Adaptation | No |
| **PFL** | Adaptation | Adaptation | No |
| **PFS** | Adaptation | Adaptation | No |
| **PSL** | Adaptation | Adaptation | No |
| **PSS** | Adaptation | Adaptation | No |
| **UFL** | Adaptation | Adaptation | No |
| **UFS** | Adaptation | No change | No |
| **USL** | Adaptation | Adaptation | No |
| **USS** | No change | Adaptation | No |

